# Living representatives of the spiralian ancestral karyotype revealed by two *Lima* genomes

**DOI:** 10.64898/2026.10.09.757954

**Authors:** France Denoeud, Benjamin Istace, Yann Dussert, Jean-Marc Aury

## Abstract

Conserved synteny enables the reconstruction of ancestral chromosome organization across deeply diverged lineages, as demonstrated across metazoans. In Spiralia, comparative genomic analyses have suggested an ancestral karyotype comprising 20 chromosomes. We report high-quality genome assemblies for two *Lima* species (*Lima caribaea* and *Lima lima*) with 20 chromosomes and show that their chromosome organization matches this ancestral state, including four previously described fusion-with-mixing events. Interestingly, other Limidae genus did not retain the ancestral karyotype: they have undergone subsequent lineage-specific fusions. The conserved organization of *Lima* chromosomes provides a rare genomic anchor for reconstructing ancestral chromosome organization and lineage-specific chromosome evolution across Spiralia and, more broadly, Metazoa.

## Introduction

Phylogenetically distant metazoan genomes display remarkably conserved synteny over hundreds of millions of years, enabling the identification of ancestral linkage groups and the reconstruction of chromosomal rearrangements that shaped extant karyotypes (Putnam et al. 2007; Putnam et al. 2008; Simakov et al. 2022; Schultz et al. 2023; Lewin, Liao, et al. 2025; Schultz et al. 2026). In particular, the ancestral genome of the spiralian lineage, including Lophotrochozoa (Mollusca, Annelida, Platyhelminthes) and Gnathifera (Laumer et al. 2015; Piovani et al. 2025), was suggested to contain 20 chromosomes, resulting from four fusions relative to the bilaterian ancestral genome (N=24) (Simakov et al. 2022; Martín-Zamora et al. 2023; Song et al. 2023; Lin et al. 2024; Piovani et al. 2025; Sigwart et al. 2025; Struck 2026). Although additional genomes of Spiralia outside Lophotrochozoa are needed to more definitively establish the karyotype of the ancestral Spiralia, we hereafter refer to the 20 previously described ancestral linkage groups (ALG) as spiralian ALGs for consistency with previous literature.

The spiralian reference genomes most commonly used for metazoan synteny studies are those of scallops (Bivalvia, Pectinidae), *Patinopecten yessoensis* (Wang et al. 2017; Simakov et al. 2022) and *Pecten maximus* (Kenny et al. 2020; Lewin, Liao, et al. 2025). These genomes have 19 chromosomes and share the same fusion between two bilaterian ALGs. Chromosome-scale assemblies from several other bivalve lineages have also revealed a haploid chromosome number of 19 (Ran et al. 2019; Sun et al. 2022; Law et al. 2025). However, comparative synteny analyses reveal that this shared chromosome number arose through different lineage-specific fusion events (Lin et al. 2024; Istace et al. 2026).

Large sequencing projects aiming to produce chromosome-scale reference genomes provide unprecedented resources for studying the biology and evolution of marine organisms (The Darwin Tree of Life Project Consortium 2022; Blaxter et al. 2025; Formenti et al. 2026). As part of the ATLASea programme (https://www.atlasea.fr/), chromosome-scale genomes have recently been generated for two bivalve species of the genus *Lima*, representing the first reported spiralian genomes exhibiting a 20-chromosome karyotype. The family Limidae comprises bivalves with a semi-mobile lifestyle, combining swimming with attachment to the substrate by byssal threads, and distinctive morphological traits such as well-developed eyes (Mikkelsen and Bieler 2003; McElroy et al. 2024).

Here, we compared the two *Lima* genomes with other spiralian genomes to determine whether their 20-chromosome karyotype corresponds to the previously proposed ancestral spiralian chromosome organization. We find that *Lima* species retain this organization, including the four previously described fusion-with-mixing events, providing a valuable reference for investigating the evolution of spiralian karyotypes.

## Results

As of October 1, 2026 a survey of chromosome-scale spiralian genome assemblies available in the NCBI Genome database (https://www.ncbi.nlm.nih.gov/datasets/genome/) identified 506 reference chromosome-scale genomes assemblies, of which only two have 20 chromosomes (excluding one hybrid between two 10 chromosome species). These two species belong to the genus *Lima* (Mollusca/Bivalvia/Limida/Limidae): *Lima caribaea* (GCA_977909015.1) and *Lima lima* (GCA_984582965.1). Both assemblies were generated using PacBio HiFi and Hi-C data by the ATLASea consortium, and they meet the Earth BioGenome Project reference genome quality standards (Lawniczak et al. 2022). The Hi-C contact maps were used to validate the chromosome-scale structure of both assemblies. They showed no structural ambiguities and supported a karyotype of 20 chromosomes for both *L. lima* and *L. caribaea* (Fig. S1).

To investigate synteny conservation relative to the ancestral spiralian structure, we ran Hobrac (Istace et al. 2026 ; https://github.com/Genoscope-LBGB/hobrac) on six Limidae species and two 19-chromosome bivalves: *Pecten maximus* and *Donax variegatus* (Table S1). The Hobrac software allows to visualize synteny using mollusca_odb12 BUSCO (Manni et al. 2021; Tegenfeldt et al. 2025) genes, detect significant chromosome associations, and color them according to the 24 corresponding bilaterian ALGs described previously (Simakov et al. 2022). The synteny plots confirm that the 20 *Lima* chromosomes correspond to the 20 previously described spiralian ALGs, with the 4 fusion-with-mixing events, L□J2, H□Q, K□O2, and O1□R, already documented (Simakov et al. 2022; Lin et al. 2024; Struck 2026) (Fig. 1). In addition to the shared four ancestral events, the comparison reveals lineage-specific fusions-with-mixing. Among Limidae, two fusions are shared by the two *Ctenoides* species and a distinct set of four fusions are shared by the two *Limaria* species, with two additional fissions possibly having occurred in *L. hians*. Inspection of the Hi-C contact map does not provide sufficient evidence to confidently establish whether these apparent splits are genuine or assembly artefacts (Fig. S2). Dot plots further reveal detectable microsynteny within each pair of closely related species, whereas gene order is substantially less conserved across genera (Fig. 2). The macrosynteny rearrangement indices (RI) (Lewin et al. 2024) relative to the 24 bilaterian ALGs were calculated with Hobrac. As expected, both *Lima* species have lower RI than the other bivalve species considered, with *Lima caribaea* showing the lowest index and thus the closest overall match to the ancestral chromosome organization.

**Fig. 1.**
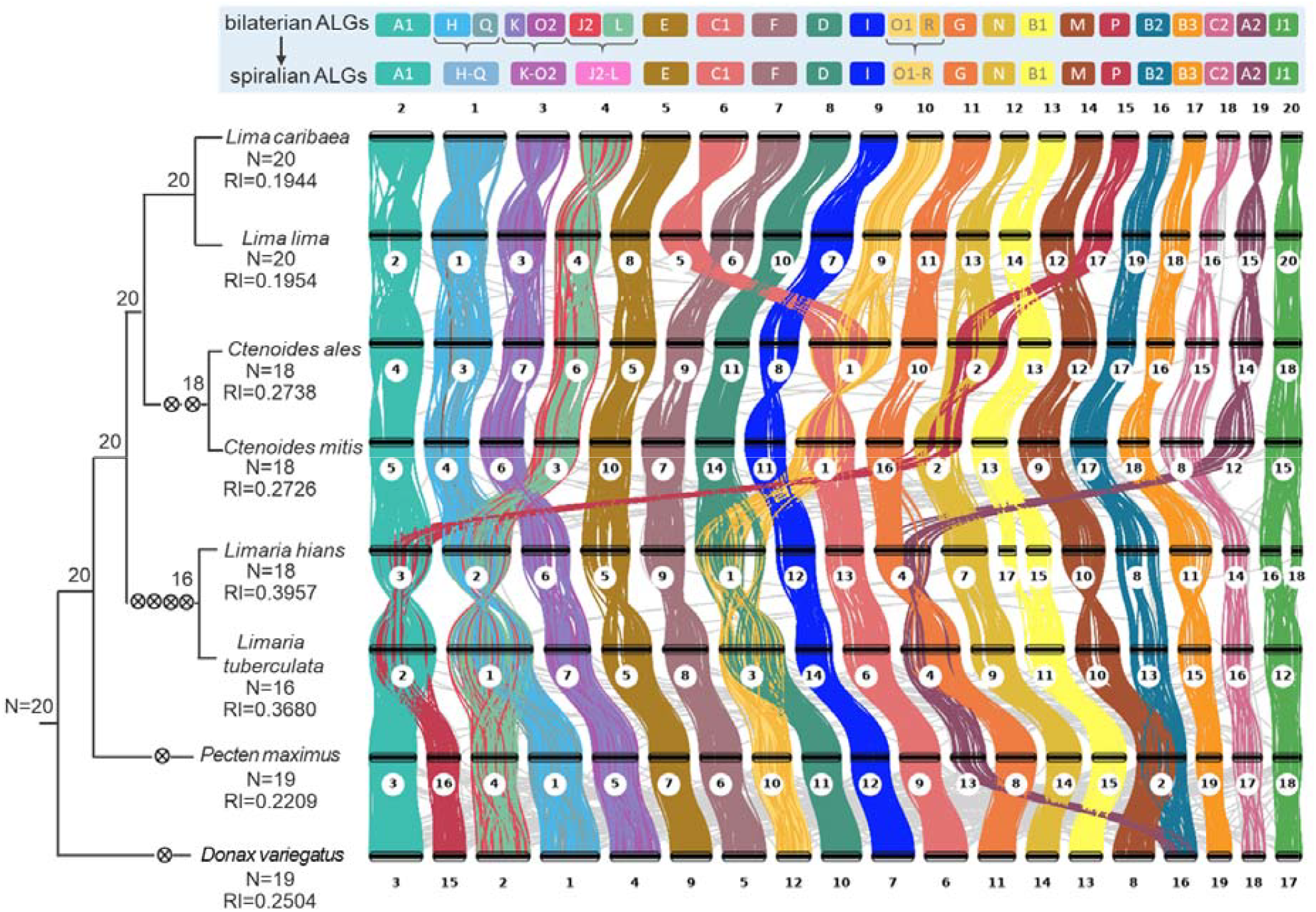
Ribbon plot displaying syntenic relationships between eight bivalve genomes. Syntenic relationships are based on mollusca_odb12 BUSCO genes and were generated with Hobrac. BUSCO genes are colored according to the previously described 24 bilaterian ancestral linkage groups (ALGs), but only genes belonging to valid chromosome chains are colored (N=3051). The upper panel shows the fusion-with-mixing events leading from the 24 bilaterian ALGs to the 20 spiralian ALGs. The phylogeny, represented by a cladogram, was reconstructed using maximum likelihood inference from the sequences of single-copy BUSCO proteins predicted in the 8 species. All branches have 100% bootstrap support. Fusion-with-mixing events are indicated by ⊗. Rearrangement index (RI) values were computed with Hobrac. The number of chromosomes and RI are displayed under each species name.

**Fig. 2.**
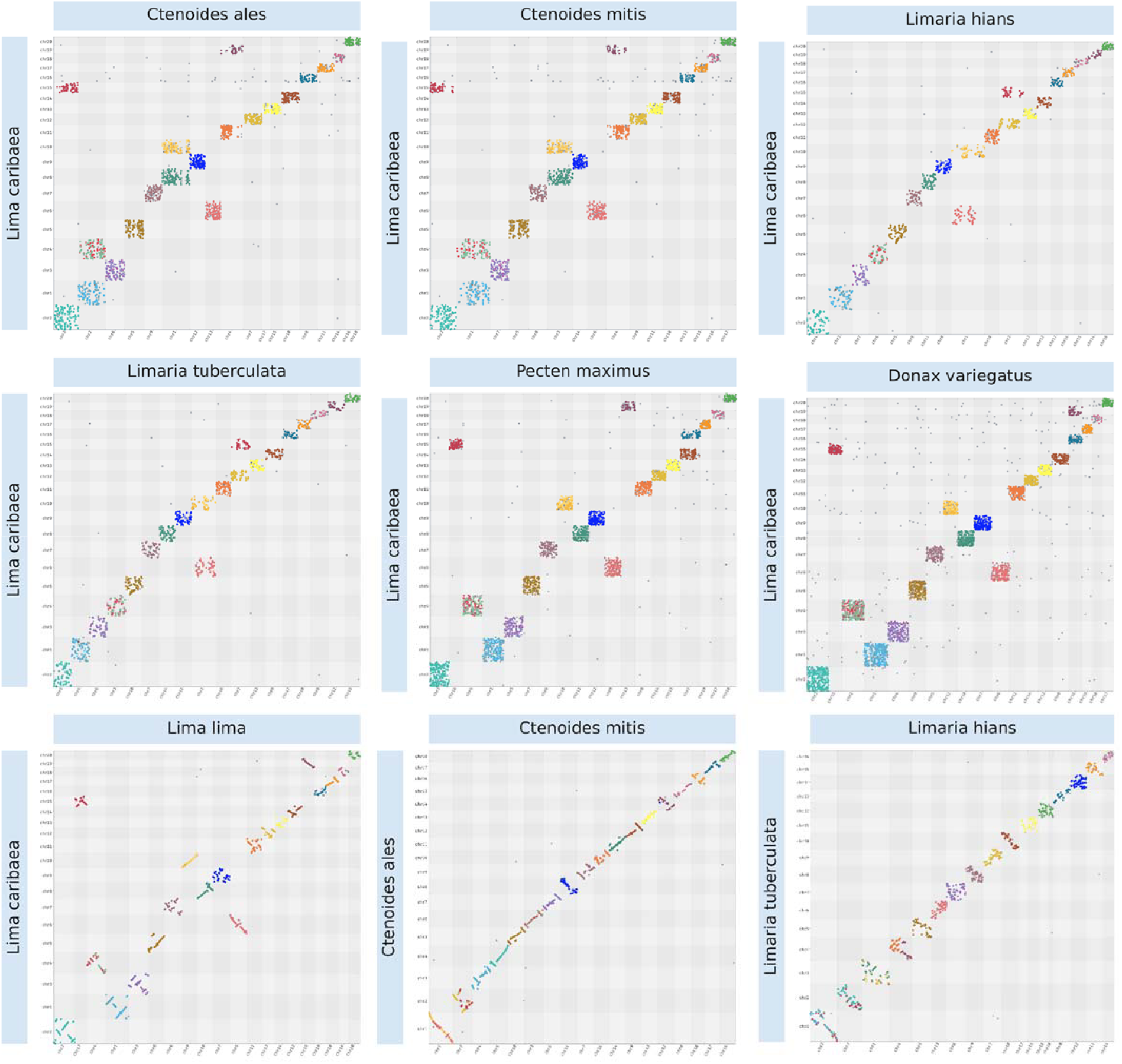
Dotplots displaying syntenic relationships between eight bivalve genomes. The first seven dot plots were generated with Hobrac using the seven genomes displayed in Fig. 1, with each dot representing the chromosomal location of a *mollusca_odb12* BUSCO gene in the two compared genomes. The genome of *Lima caribaea* is compared with the seven other genomes. Two additional Hobrac runs were performed to generate dotplots comparing the two *Limaria* species and the two *Ctenoides* species.

To illustrate the utility of *Lima caribaea* as a reference genome for synteny analyses, we also compared *Lima caribaea* with five molluscs outside Bivalvia (*Pictodentalium vernedei* (Song et al. 2023), *Bolma rugosa, Acanthopleura gemmata, Deshayesiella sirenkoi* (Sigwart et al. 2025), *Chaetoderma sp* (Wang et al. 2024)), five lophotrochozoans outside molluscs (*Urechis unicinctus* (Cheng et al. 2024), *Arenicola marina, Lineus longissimus, Discradisca antillarum* (Roberts et al. 2025), *Phoronis australis* (Lewin, Sakagami, et al. 2025)), and one non-parasitic platyhelminthes, *Pseudobiceros splendidus* (Table S1). The synteny plots confirm that all the compared Lophotrochozoa share the four previously described fusion-with-mixing events, suggesting that their common ancestor likely had 20 chromosomes. The four ancestral fusions are also shared by *P. splendidus*, although subsequent fusions followed by a probable whole genome duplication event complicate the pattern (Fig. 3, Fig. S3).

**Fig. 3.**
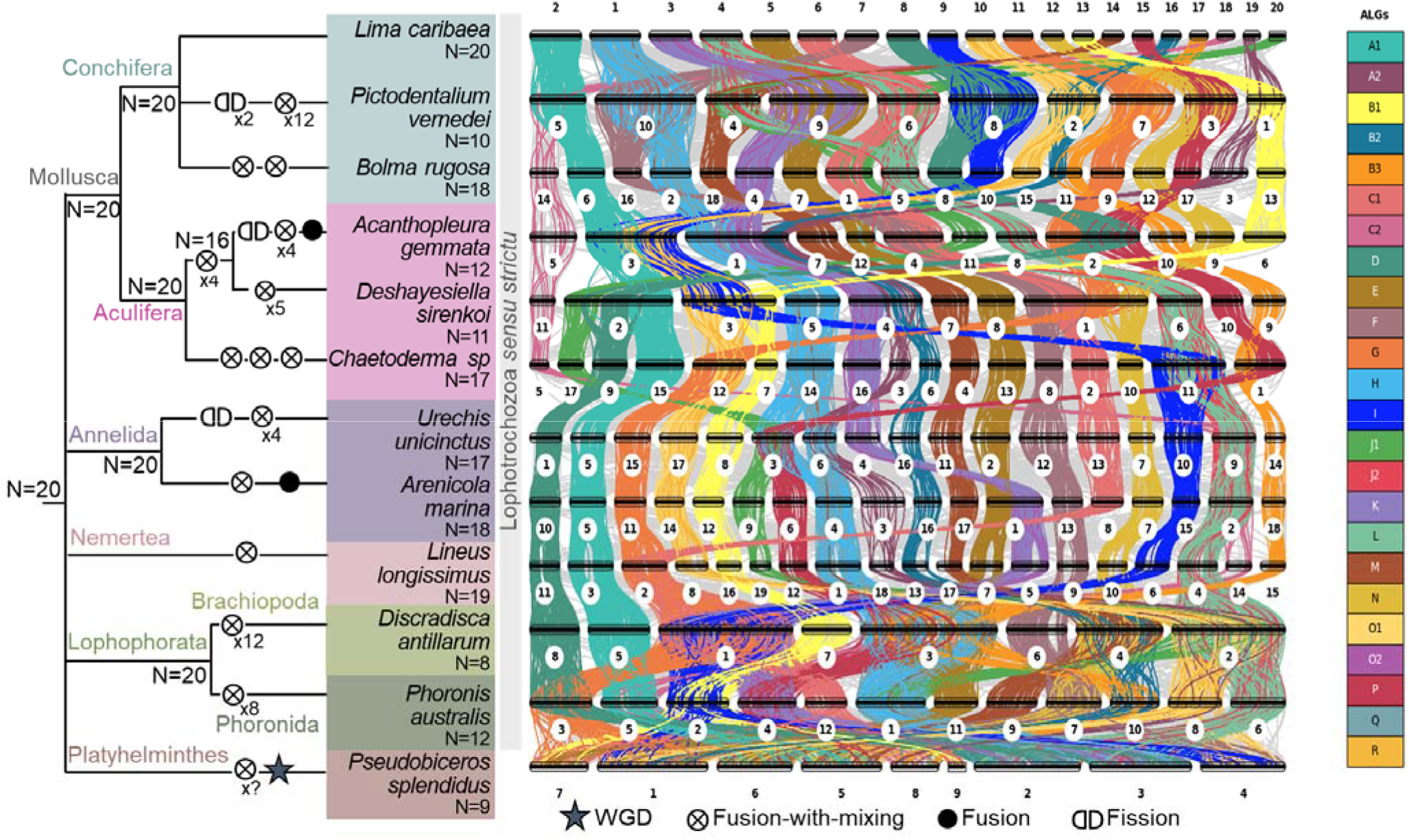
Syntenic relationships across 12 spiralian genomes. The ribbon plots were generated with Hobrac. The BUSCO *mollusca_odb12* genes are colored according to the previously described 24 bilaterian ALGs, but only genes belonging to valid chromosome chains are colored. The phylogeny was adapted from recent studies (Chen et al. 2025; Struck 2026), with polytomies where relationships between lineages remain unresolved. The putative history of chromosomal rearrangements leading to the karyotypes of extant species was inferred from the ribbon plots and associated dotplot grids (Fig. S3). The rearrangement events were placed arbitrarily along the branches and are not intended to represent a temporal scale.

Only three chromosome-scale genome assemblies of spiralian species outside Lophotrochozoa+Platyhelminthes are currently available, in the Gnathifera lineage. One is a rotifer, *Adineta vaga*, that displays extensive chromosomal scrambling (Flot et al. 2013; Simion et al. 2021). The two others are the chaetognaths *Flaccisagitta enflata* (Guiglielmoni et al. 2025) and *Paraspadella gotoi* (Piovani et al. 2025), that both contain 9 chromosomes. *Paraspadella gotoi* was shown to retain remnants of the ancestral bilaterian ALGs, although only two of the four ancestral fusions could be identified and some of the ALGs involved could not be recovered (Piovani et al. 2025; Struck 2026). Comparing *Lima caribaea* and the two chaetognath genomes revealed a one-to-one correspondence between *P. gotoi* and *F. enflata* chromosomes, and confirmed previous findings by detecting only seven significant (P ≤ 0.05) chromosome associations shared with *Lima caribaea*, two of which correspond to the ancestral fusions K⊗O2 and O1⊗R (Fig. S4).

Finally, we included the assembly of *Lima caribaea* to generate a reference mapping between BUSCO gene sets and the 20 spiralian ancestral linkage groups, based on conserved chromosomal locations across seven metazoan species following a previously described approach (Istace et al. 2026). This mapping was used to generate ALG-based color files for BUSCO datasets, made available for use with Hobrac (option --color-spiralian-alg). It provides a reusable resource for the community, enabling other researchers to investigate chromosome structure evolution across spiralian lineages without reconstructing ALG assignments.

Of the 4,281 complete single-copy *mollusca_odb12* BUSCO genes detected in *Lima caribaea*, 3,899 (91.1%) were assigned to a spiralian ALG. Among these, 3,872 (99.3%) were located on the *Lima* chromosome corresponding to the assigned ALG. Thus, 90.4% of the detected BUSCO genes were both assigned to an ALG and located on the corresponding *Lima* chromosome, supporting the use of *Lima* chromosome locations as a reliable proxy for ancestral spiralian ALG organization. This chromosome-based proxy can then be used to extend ALG-based analyses to orthologous gene sets beyond BUSCO without requiring a new ALG mapping.

## Discussion

Our analyses identify *Lima caribaea* and *Lima lima* as the first chromosome-scale spiralian genomes known to retain the 20 ancestral linkage groups previously proposed for the spiralian ancestor. Their conserved chromosomal organization, including the four ancestral fusion-with-mixing events, supports the proposed 20-chromosome ancestral organization within the molluscan and lophotrochozoan lineages examined here, while additional chromosome-scale genomes from other spiralian lineages will be needed to determine whether this organization extends more broadly across Spiralia.

Within Limidae, *Lima* retains the proposed ancestral organization, whereas *Ctenoides* and *Limaria* show distinct additional fusions, indicating independent chromosome evolutionary histories. Thus, the ancestral organization observed in *Lima* occurs within a family that has experienced substantial chromosomal diversification.

The conserved chromosome-ALG associations also provide a practical framework for comparative genomics. In particular, the low rearrangement index of *Lima caribaea* makes it a valuable reference genome for investigating chromosome evolution, while its conserved chromosome-ALG associations provide a direct framework for visualizing ancestral linkage groups directly on synteny plots. These conserved associations can also be used to extend ALG assignments to orthologous gene sets beyond BUSCO genes. As chromosome-scale genomes accumulate across Spiralia, *Lima* therefore provides a genomic anchor against which ancestral chromosomal features and lineage-specific rearrangements can be disentangled.

## Methods

### Data retrieval

All genome assemblies used in this study were retrieved using ncbi-datasets (O’Leary et al. 2024) with the accession numbers listed in Table S1.

### Hi-C contact map generation

Long-reads and Hi-C sequencing data as well as reference chromosomes were downloaded from the ENA (PRJEB111915 for *Lima lima*, PRJEB97432 for *Lima caribaea* and PRJEB75297 for *Limaria hians*). Hi-C reads were aligned to the chromosomes following the Arima mapping pipeline and the resulting alignments were used to generate the contact map with PretextMap v1.0.0 (Howe et al. 2021) with the following arguments: “--highRes --sortby nosort --mapq 0”. Long reads were then aligned to the chromosomes using Minimap2 (Li 2018) and only primary alignments were retained by using samtools (Li et al. 2009) view -F 2308. These long-reads alignments were then given as input to PretextMap in order to generate the coverage track of the contact maps (Fig. S1-2). The Hi-C contact maps are available at https://zenodo.org/records/23013306.

### Synteny analyses

Hobrac was run on three distinct datasets: 8 bivalves (Fig. 1-2), 11 lophotrochozoans + the platyhelminth *P. splendidus* (Fig. 3, S3) and *Lima* + 2 chaetognaths (Fig. S4). All runs were performed with the following options : --color-bilaterian-alg --min-busco-genes 10 --alg-pvalue 0.05. Conserved chromosome chains were detected as described in (Istace et al. 2026). Briefly, significant associations between chromosome pairs were identified using Fisher’s exact tests with Bonferroni correction, using a corrected significance threshold of 0.05. Significant chromosome associations were then connected across species to identify conserved chromosome chains, which were filtered and validated according to the criteria described in (Istace et al. 2026). For visualization, genes belonging to chromosome chains retained after validation were colored using the precomputed file assigning each BUSCO gene to its corresponding bilaterian ALG.

### Phylogeny reconstruction in Limidae and two outgroup bivalves

The genomes of the six Limidae species and of *Pecten maximus* and *Donax variegatus* were analyzed with BUSCO 6.0.0 (--metaeuk parameter) with the mollusca_odb12 database. We selected the single copy BUSCOs present in all species, which represented 3,047 genes. For each gene, multiple alignments of protein sequences were produced with MAFFT 7.525 (-- auto --maxiterate 1000) (Katoh and Standley 2013) and cleaned with trimAl 1.5.0 (- automated1) (Capella-Gutiérrez et al. 2009). After concatenation of the alignments into a supermatrix, maximum likelihood phylogenetic reconstruction was carried out with IQ-TREE 3.0.1 (Wong et al. 2026), using inference of the best-fitting partition and substitution models (Chernomor et al. 2016; Kalyaanamoorthy et al. 2017) and assessing branch support with 1,000 ultrafast bootstrap replications (Hoang et al. 2018). The phylogeny obtained is concordant with previous literature (Dougherty and Li 2017).

### Assignment of BUSCO genes to spiralian ALGs

We generated a mapping of BUSCO genes to the 20 spiralian ALGs following the procedure described for Hobrac, with *Lima caribaea* added as a seventh reference genome. The correspondence between conserved chromosome chains and the previously defined ALGs was established from synteny analyses of seven chromosome-scale genomes (*Branchiostoma floridae, Holothuria leucospilota, Pecten maximus, Donax variegatus, Nematostella vectensis, Rhopilema esculentum*, and *Lima caribaea*). A BUSCO gene was assigned to an ALG when it was detected on the corresponding chromosome in at least three genomes, with one discordant chromosome allowed when the gene was detected in at least five genomes (except *B. floridae* for the R group). The resulting mapping was used to generate ALG-based color files for various BUSCO odb12 datasets and is provided with the Hobrac color files (https://github.com/Genoscope-LBGB/hobrac/tree/master/hobrac/colors).

### Evaluation of *Lima caribaea* chromosomes as proxies for spiralian ALGs

To evaluate the correspondence between *Lima caribaea* chromosomes and the 20 spiralian ALGs, we considered all mollusca_odb12 BUSCO genes detected (S: complete and single-copy) in the *Lima caribaea* assembly. For each gene assigned to a spiralian ALG by the reference mapping, we compared the chromosome carrying the gene in *Lima caribaea* with the chromosome corresponding to the assigned ALG. Genes located on the expected chromosome were considered concordant, whereas genes located on a different chromosome were considered discordant. BUSCO genes not assigned to an ALG by the reference mapping were excluded from this concordance assessment.

A chromosome-based color file was generated for *Lima caribaea* by assigning each detected BUSCO gene the color corresponding to its chromosome (Table S2). It can be provided to Hobrac with the option --custom-colors.

## Supplementary material

Supplementary_file_1.pdf: contains Supplementary Table S1 and Supplementary Figures S1-4.

Supplementary_table_2.xlsx

## Data availability

The genome assemblies used during the current study are available in the NCBI under the accession numbers listed in Table S1. Pretext Hi-C contact maps generated for *Lima caribaea, Lima lima* and *Limaria hians* are available at https://zenodo.org/records/23013306. Hobrac is an open source software, source code and binaries are freely available from https://github.com/Genoscope-LBGB/hobrac.

## Acknowledgements

The authors acknowledge the use of AI (ChatGPT) in some sections of the manuscript to improve the English.

## Funding

This work was supported by the Genoscope, the Commissariat à l’Énergie Atomique et aux Énergies Alternatives (CEA), France Génomique (ANR-10-INBS-09-08), and the exploratory research programme ‘ATLASea: Atlas of marine genomes and its targeted projects’ WHEEL-Sea (ANR-22-EXAT-0001), DIVE-Sea (ANR-22-EXAT-0002), SEQ-Sea (ANR-22-EXAT-0003) and BYTE-Sea (ANR-22-EXAT-0004).

## Authors’ contributions

FD performed the analyses and wrote the article. BI implemented the option to use spiralian ALGs in Hobrac. YD constructed the phylogenetic tree between the six Limidae and two bivalve outgroups. JMA performed the validation of the chromosome structures with Hi-C data. BI, YD and JMA reviewed drafts of the article. All authors read and approved the final manuscript.

## Notes

https://zenodo.org/records/23013306

### Competing Interest Statement

The authors have declared no competing interest.

