## Supplementary file 1 (Fig. S1-4, Table S1) for "Living representatives of the spiralian ancestral karyotype revealed by two *Lima* genomes"

**Table S1: List of genome assemblies studied**

| Species | Lineage | Assembly AC | Chromosome number | Genome paper reference | Used for synteny plot(s): |
|---|---|---|---|---|---|
| *Lima caribaea* | Mollusca/Bivalvia | GCA_977909015.1 | 20 | | Figures 1, 2, 3, S3, S4 |
| *Lima lima* | Mollusca/Bivalvia | GCA_984582965.1 | 20 | | Figures 1, 2 |
| *Limaria hians* | Mollusca/Bivalvia | GCA_965230825.1 | 18 | | Figures 1, 2 |
| *Limaria tuberculata* | Mollusca/Bivalvia | GCA_986922335.1 | 16 | | Figures 1, 2 |
| *Ctenoides ales* | Mollusca/Bivalvia | GCA_042257175.1 | 18 | McElroy, 2024 | Figures 1, 2 |
| *Ctenoides mitis* | Mollusca/Bivalvia | GCA_978015095.1 | 18 | | Figures 1 ,2 |
| *Pecten maximus* | Mollusca/Bivalvia | GCA_902652985.1 | 19 | Kenny, 2020 | Figures 1, 2 |
| *Donax variegatus* | Mollusca/Bivalvia | GCA_964468855.2 | 19 | | Figures 1, 2 |
| *Pictodentalium vernedei* | Mollusca/Scaphopoda | GCA_031216915.1 | 10 | Song, 2023 | Figures 3, S3 |
| *Bolma rugosa* | Mollusca/Gastropoda | GCA_964656265.1 | 18 | | Figures 3, S3 |
| *Acanthopleura gemmata* | Mollusca/Polyplacophora | GCA_977063515.1 | 12 | | Figures 3, S3 |
| *Deshayesiella sirenkoi* | Mollusca/Polyplacophora | GCA_045838535.1 | 11 | Sigwart, 2025 | Figures 3, S3 |
| *Chaetoderma sp* | Mollusca/Aplacophora | GCA_034401795.1 | 17 | Wang, 2024 | Figures 3, S3 |
| *Urechis unicinctus* | Annelida | GCA_034190875.2 | 17 | Cheng, 2024 | Figures 3, S3 |
| *Arenicola marina* | Annelida | GCA_964019135.1 | 18 | | Figures 3, S3 |
| *Lineus longissimus* | Nemertea | GCA_910592395.2 | 19 | | Figures 3, S3 |
| *Discradisca antillarum* | Brachiopoda | GCA_057556925.1 | 8 | Roberts, 2025 | Figures 3, S3 |
| *Phoronis australis* | Phoronida | GCA_055505105.1 | 12 | Lewin, 2025 | Figures 3, S3 |
| *Pseudobiceros splendidus* | Plathyhelminthes | GCA_965235105.1 | 9 | | Figures 3, S3 |
| *Flaccisagitta enflata* | Chaetognatha | GCA_965153355.1 | 9 | Guiglielmoni, 2025 | Figure S4 |
| *Paraspadella gotoi* | Chaetognatha | GCA_051294375.1 | 9 | Piovani, 2025 | Figure S4 |

**Fig. S1. Hi-C contact maps of the (a) *Lima caribaea* and (b) *Lima lima* genome assemblies, generated with PretextMap.** The maps display Hi-C interaction frequencies between genomic regions, ranging from no or few interactions (light blue) to a high number of contacts (red). Chromosome-scale scaffolds are delineated by black lines and highlighted. In both species, the contact maps support the organization of the assemblies into 20 chromosomes, with no apparent structural ambiguities.

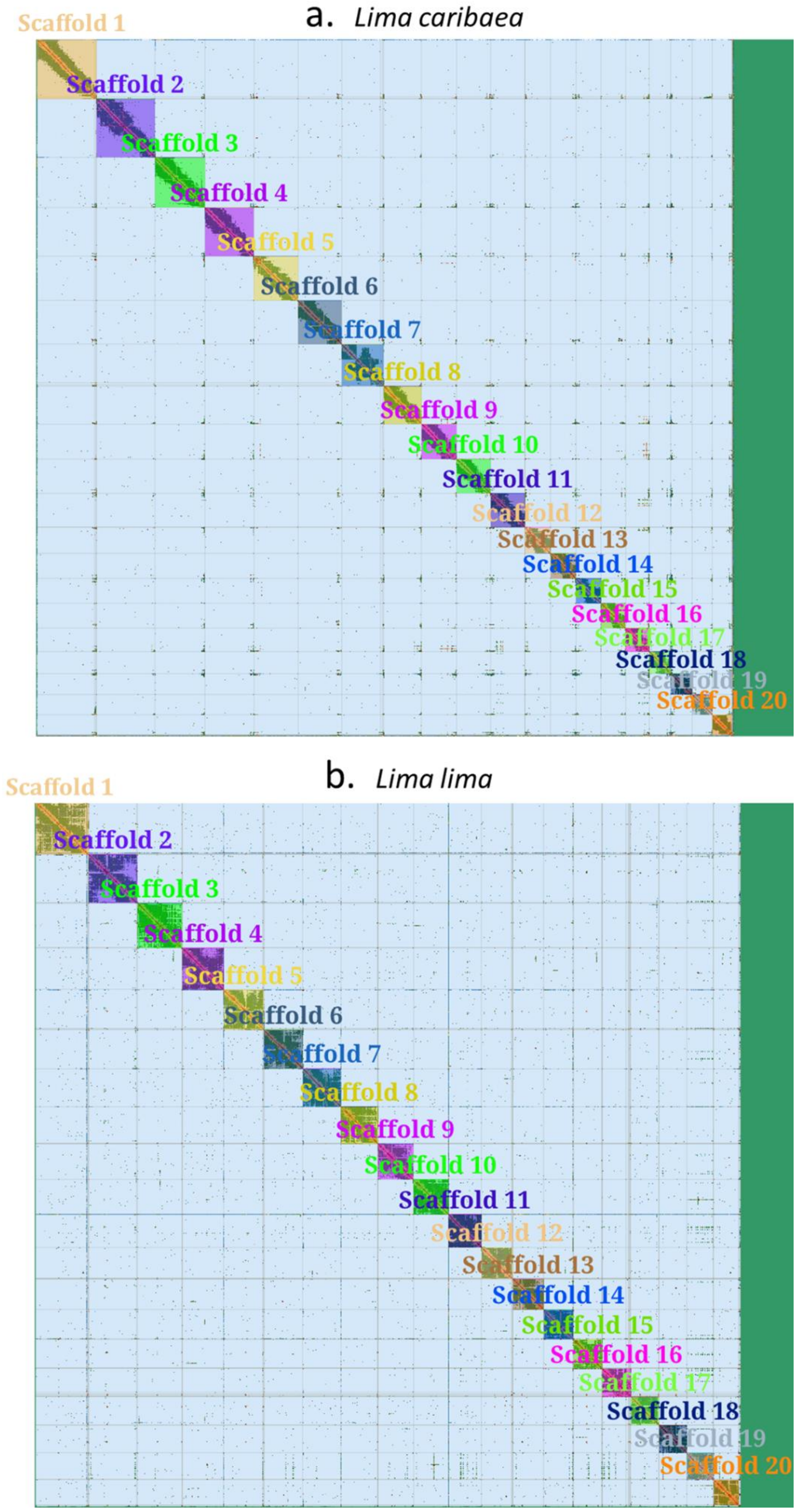

**Fig. S2. Hi-C contact map of the *Limaria hians* assembly, generated with PretextMap.** The map displays Hi-C interaction frequencies between genomic regions, ranging from no or few interactions (light blue) to a high number of contacts (red). The chromosome-scale scaffolds are delineated by black lines and highlighted, with a zoom on chromosomes 15, 16, 17, and 18. The contact map does not provide evidence to join chr15 with chr17 or chr16 with chr18.

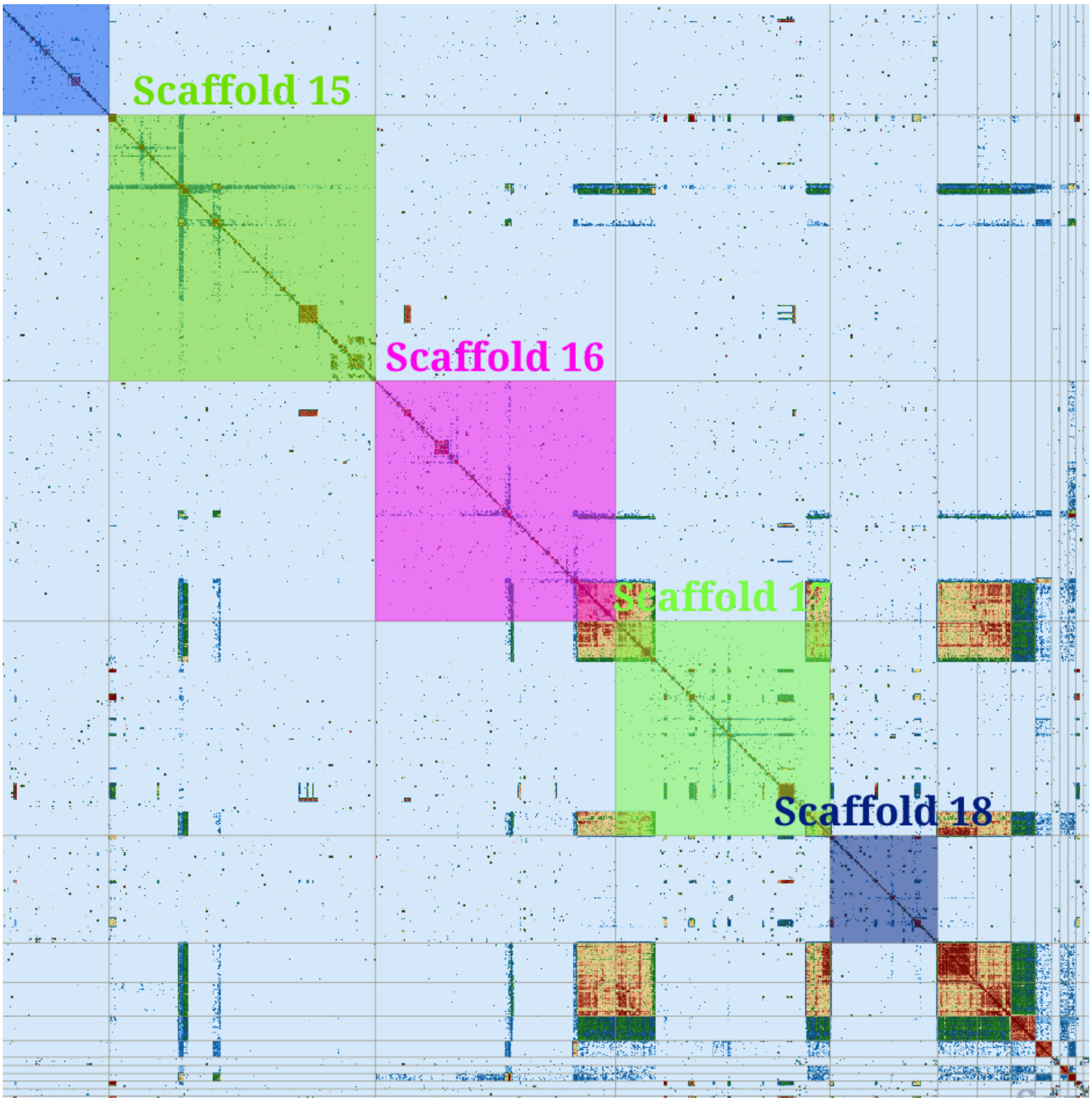

**Fig. S3. Hobrac pairwise comparison of Lima caribaea with the other 11 spiralian genomes displayed in Fig. 3.** Dots represent the chromosomal positions of BUSCO *mollusca_odb12* genes in each pair of genomes compared. Colors correspond to the 24 bilaterian ALGs. Genes that do not belong to valid chromosome chains detected by Hobrac are shown in grey.

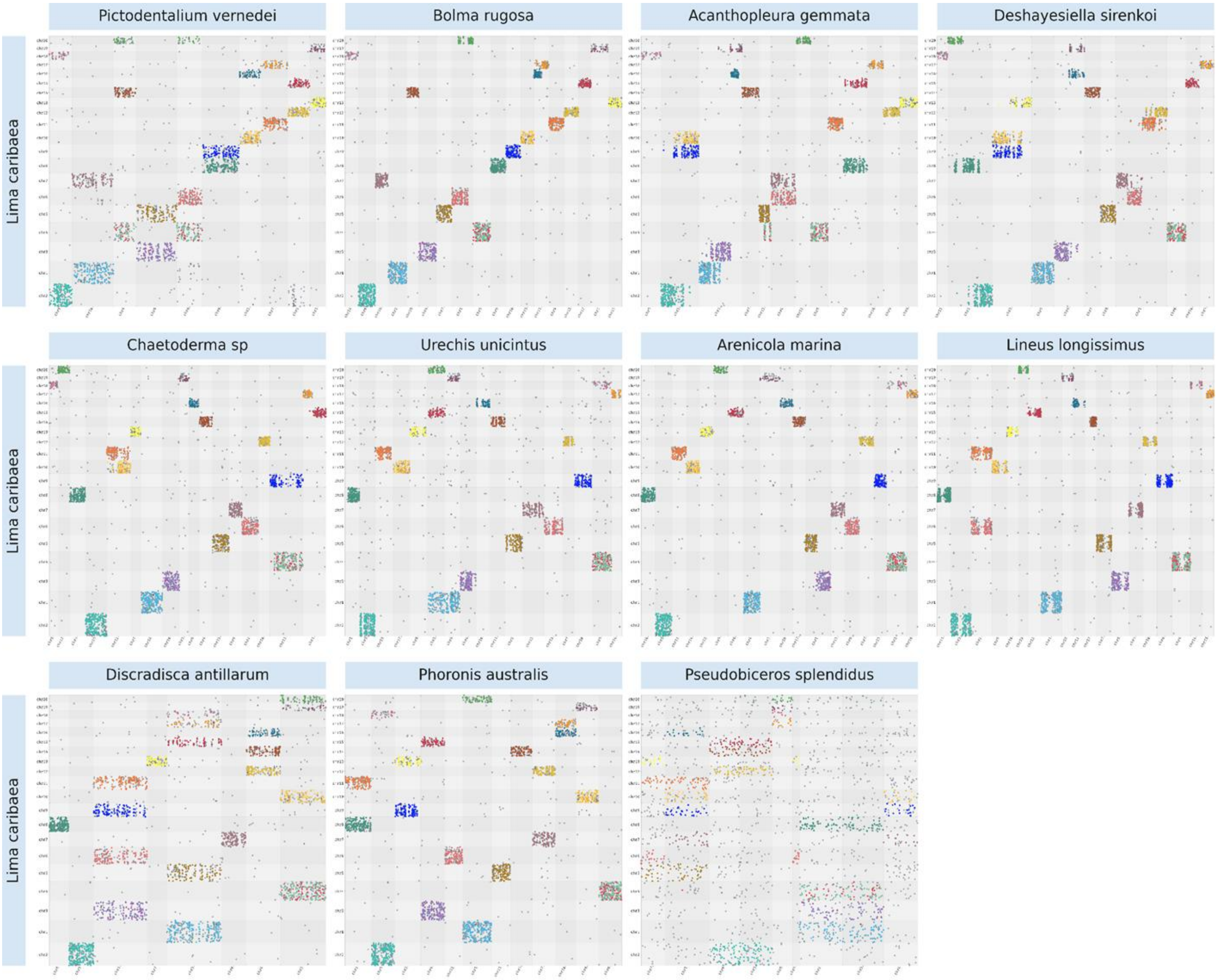

**Fig. S4. Synteny analysis between *Lima caribaea* and two chaetognaths.** a. Ribbon plot with de novo detected chromosome chains colored by default (one color for each valid chromosome association detected). b. Ribbon plot with the detected chromosome chains colored according to the mapping of BUSCO genes to bilaterian ALGs. c. Dotplot between BUSCO genes in *P. gotoi* and *F. enflata*, colored by default. d. Dotplots comparing *Lima caribaea* with the two chaetognaths, colored according to bilaterian ALGs; only the seven significant chromosome associations are colored.

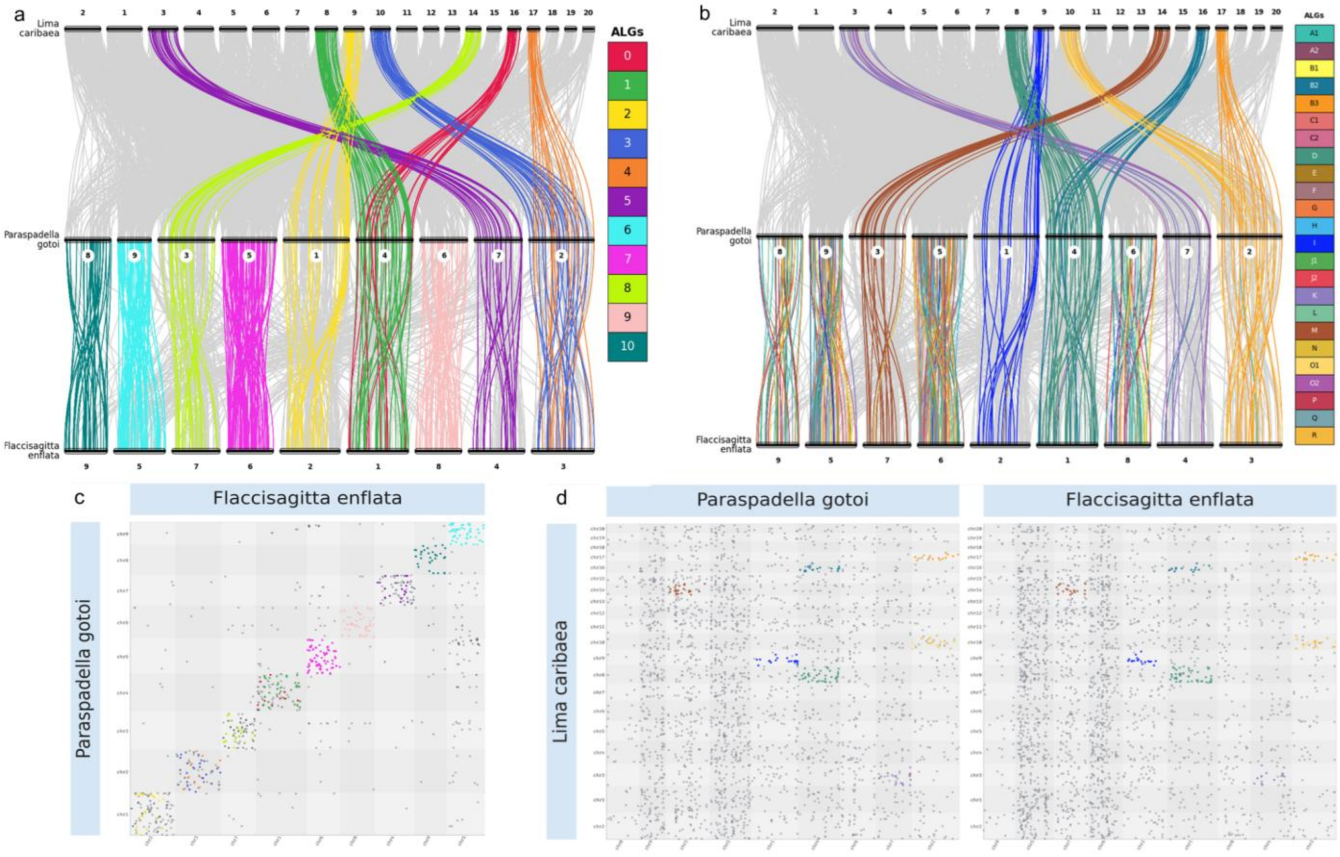
